# A senescence-like cellular response inhibits bovine ephemeral fever virus proliferation

**DOI:** 10.1101/2021.01.28.428738

**Authors:** Yu-Jing Zeng, Min-Kung Hsu, Chiao-An Tsai, Chun-Yen Chu, Hsing-Chieh Wu, Hsian-Yu Wang

**Affiliations:** International Program in Animal Vaccine Technology, International College, National Pingtung University of Science and Technology, Pingtung, Taiwan R.O.C; Research Center for Tumor Medical Science, China Medical University, Taichung, Taiwan R.O.C; Graduate Institute of Animal Vaccine Technology, College of Veterinary Medicine, National Pingtung University of Science and Technology, Pingtung, Taiwan R.O.C

**Keywords:** anti-viral response, senescence-like response, viral vaccine production, BEFV, Rhabdoviridae, GSEA, BHK-21, bioreactor

## Abstract

During industrial scale production of virus for vaccine manufacturing, antiviral response of host cells can dampen maximal viral antigen yield. In addition to interferon responses, many other cellular responses such as the AMPK signaling pathway or senescence-like response may inhibit or slow down virus amplification in the cell culture system. In this study, we first performed a Gene Set Enrichment Analysis of the whole-genome mRNA transcriptome and found a senescence-like cellular response in BHK-21 cells when infected with bovine ephemeral fever virus (BEFV). To demonstrate that this senescence-like state may reduce virus growth, BHK-21 subclones showing varying degrees of senescence-like state were infected with BEFV. Results showed the BHK-21 subclones showing high senescence staining could inhibit BEFV replication while low senescence-staining subclones are permissive to virus replication. Using a different approach, a senescence-like state was induced in BHK-21 using a small molecule, camptothecin (CPT), and BEFV susceptibility was examined. Results showed that indeed CPT-treated BHK-21 is more resistant to virus infection. Overall, these results indicate that a senescence-like response may be at play in BHK-21 upon virus infection. Furthermore, cell clone selection and modulating treatments using small molecules may be tools in countering anti-viral responses.

## Introduction

Successful virus production using cell cultures depends in large part on the response of the host cells. Some cell lines, such as CHO, are highly resistant to virus infection due to their strong anti-viral interferon response. In contrast, some cell lines, such as Vero and BHK-21, exhibit low interferon response and are therefore more amenable to virus production for purposes such as vaccine manufacturing (1–3). Upon virus infection, in addition to interferon responses, a host of other types of responses may also come into play inside the host cell: senescence-like response, the AMPK signaling pathway, or innate immune responses that can inhibit or slow down virus amplification. These anti-viral responses may be further enhanced in the large-scale bioreactor because of the multiple amplification of the production cells in the culture vessel. To understand and manage anti-viral responses of the host cell are critical steps for the development of an optimal virus culturing system.

Bovine ephemeral fever virus (BEFV) belongs to the *Rhabdoviridae* family with a single-stranded, negative sense RNA genome that encodes five structural proteins (N, P, M, G, and L) and a nonstructural proteins (4–7). Although mortality due to BEFV infection is low, reduced milk production during pandemic periods can result in significant economic losses for dairy farms (8). While inactivated BEFV vaccines provide sufficient protection, immunity decreases rapidly and cattle need to be vaccinated every one to two years (9). Therefore, a stable production of this virus as vaccine antigen is important for the cattle industry.

BHK-21 is widely used for BEFV production because of its high virus susceptibility. Clear cytopathic effect (CPE) and cell death can be observed 48 hours after BEFV infection. However, some BHK-21 cells appear not to show CPE, remain alive, and can even be passaged. The survival of these cells could mean a less efficient in virus production system, thus, it is important to explore potential explanations for the survival of some BHK-21 cells after BEFV infection. In the previous studies, not only the foot-and-mouth disease virus but also the rabies virus had reported the persistent infection in BHK-21 cells (10, 11). We also noticed that the BEFV persistent infected BHK-21 cells become enlarged and flattened morphology which similar to the senescence cells described by the other researchers (12, 13).

In this study, we first performed a cell functional analysis of BEFV-infected BHK-21 cells to look for gene sets are enriched after infection. Enriched gene sets may point to potential pathways of anti-viral response. We then hypothesized that a senescence-like state may play a role in virus-resistance of some BHK-21 cells. As supporting evidence, we performed two experiments: 1) subclones of BHK-21 were stained for senescence-like state and their resistance to BEFV infection was studied, and 2) a senescence-like state was induced in BHK-21 using a small molecule, camptothecin (CPT), and BEFV susceptibility was examined.

## Materials and methods

### BEFV culture and tittering

Baby Hamster Kidney-21 cells (BHK-21, BCRC#60041, purchased from the Bio-resource Collection and Research Center, Taiwan) were cultured in growth medium (MEM, Gibco co. LTD, USA, with 10% FBS, Invitrogen) at 37°C in an atmosphere of 5% CO_2_. The BEFV strain Tn88128 isolated in Taiwan (4) was cultured in BHK-21 cells (MOI = 0.01) in 2% FBS media (MEM media containing 2% FBS) at 37 °C with 5% CO_2_. The mock group cells were incubated with the 2% FBS media only. Virus titer was determined by the TCID_50_ or q-RTPCR method. For TCID_50_, CPE was observed under the microscope 3 days after inoculation and calculated by using the Reed and Muench method (14). For q-PCR analysis of virus titer, viral RNA was isolated with QIAzol Lysis Reagent (Qiagen Co. LTD, USA). After reverse-transcription by the SensiFAST™ cDNA Synthesis Kit (Bioline, Memphis, USA), cDNA samples were prepared with the QuantiFast SYBR Green PCR Master Mix reagent (Qiagen, Hilden, Germany) with the detection primers (BEFV-G F: 5’-TACCCTCCTGCTG-GATGCTTTTG-3’ and BEFV-G R: 5’-CTGTGTGCATTCTAAAACCTGGC-3’). Real-time PCR conditions were 95°C for 3 min, followed by 40 cycles at 95°C for 15 s, 60°C for 30 s and 72°C for 15 s. The virus titer was determined by comparing to the standard virus stock and shown as TCID_50_/ml.

### Cell functional analysis for surviving BHK-21 cells after BEFV infection (Gene Set Enrichment Analysis, GSEA)

BHK-21 cells were inoculated with BEFV and total cellular mRNA were harvest at the indicated time with the QIAzol Lysis Reagent (Qiagen Co. LTD, USA) according to the protocol of the manufacturer. Total cDNA was prepared by the SensiFAST™ cDNA Synthesis Kit (Bioline, Memphis, USA) and analyzed by next generation sequencing service (Illumina, 150pb pair end, 6G, Tools co. LTD. Taiwan). All NGS samples were aligned to the BHK-21 genome (Mesocricetus_auratus. MesAur 1.0.98) by using of the STAR aligner (15, 16). The gene expression were determined as FPKM by using the Cufflinks (17) to assemble and quantify transcripts FPKM. To evaluate biological pathways altered between BEFV-infected BHK-21 cell and mock-infected BHK-21, the Gene Set Enrichment Analysis (GSEA) (Broad Institute, Cambridge, MA, java GSEA Desktop Application version 4.1.0) (18, 19) tool was employed. The KEGG pathway gene subsets of GSEA Molecular Signatures Database (MSigDB) (18) C2-curated gene sets (186 gene sets) along with external senescence-related gene subsets, were applied to enrichment analysis (total 204 gene sets). The external senescence-related gene subsets were added in the analysis as the references described in **Supplementary table 1**. Gene set enrichment was considered significant at FDR< 0.25 (18).

**Table 1.**
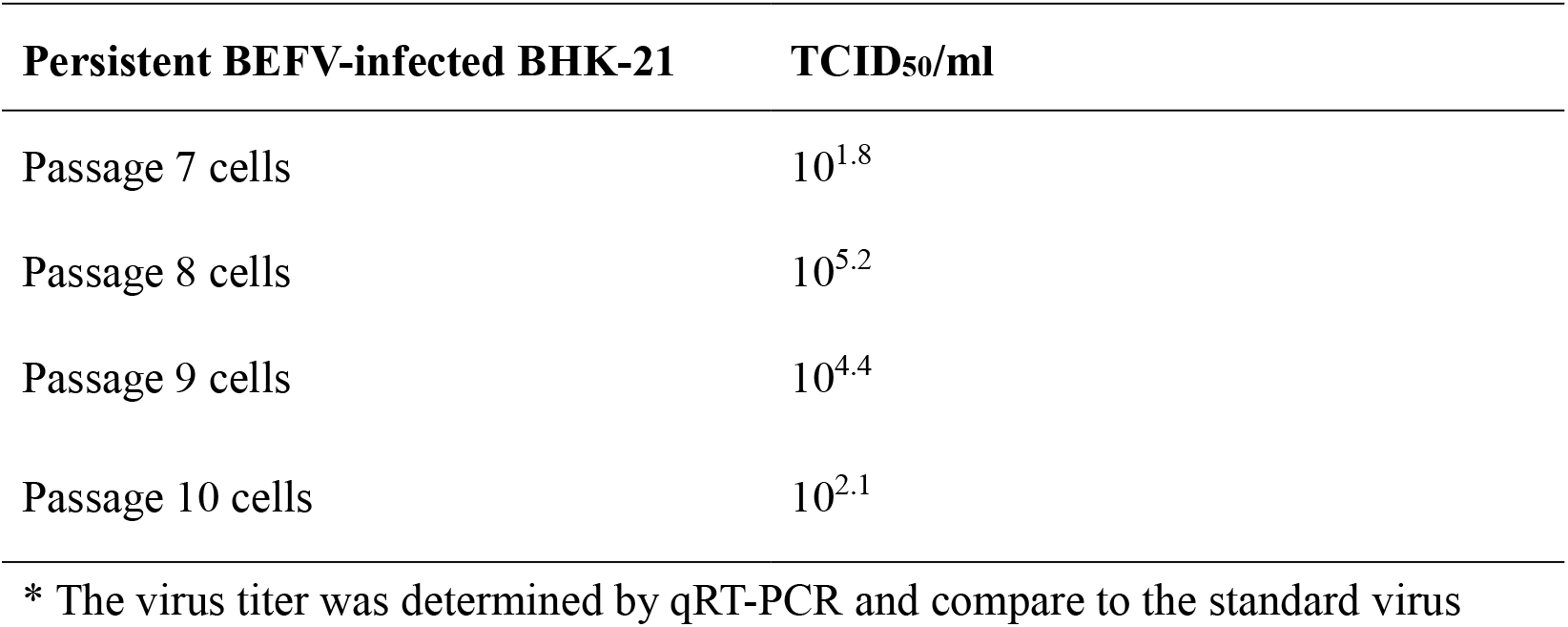
Virus titer of the persistent BEFV-infected BHK-21 cells

| **Persistent BEFV-infected BHK-21** | **TCID<sub>50</sub>/ml** |
| --- | --- |
| Passage 7 cells | 10<sup>1.8</sup> |
| Passage 8 cells | 10<sup>5.2</sup> |
| Passage 9 cells | 10<sup>4.4</sup> |
| Passage 10 cells | 10<sup>2.1</sup> |

\* The virus titer was determined by qRT-PCR and compare to the standard virus

### BHK-21 subclones isolation

To isolate subclones of BHK-21, parental BHK-21 cells were trypsinized and resuspended in growth media for calculation. The resuspended cells were diluted into 1 cell/0.1ml density and seeded into 96-well plates (corning, co. LTD, USA) (0.1 ml/well). Confluent cells were then subcultured into 24-well and 6-well plates (corning, co. LTD, USA) sequentially. Subclones were then stored in freezing media (growth media with 7% DMSO) at −80°C for further use.

### Senescence-like-associated SA-β-galactosidase staining

SA-β-gal staining was chosen as the senescence marker (20) and performed according to a protocol published by Dimri *et al.* 1995 (21). Cells were washed twice with PBS and fixed by 4% paraformaldehyde for 5 min at room temperature and washed with PBS twice. The fixed cells were stained with the SA-β-Gal staining solution (1 mg/ml X-Gal in dimethylformamide, pH 6.0 citric acid/sodium phosphate buffer, 5mM potassium ferrocyanide, 5 mM potassium ferricyanide, 150 mM NaCl, 2mM MgCl_2_) and incubated at 37°C for 12~16 hours.

### Dynamic monitoring of cell adhesion by iCELLigence™

The RTCA iCELLigence™ system (ACEA Bioscience, Inc., San Diego, CA, USA) records impedance in real-time and converts into cell index (CI) (22). This can quickly analyze cell adhesion ability and cell size during cell proliferation. The LS-BHK-21, BHK-21-parental and HS-BHK-21 cells were loaded into E-Plate L8 (ACEA Biosciences, Inc.) at 2×10^4^ cells/well and the CI was recorded at 5-minute intervals for the first 12 hours and then at 15-minute intervals till the end of the measurement period. This is convenient to monitor the cell adhesion ability and cell events during the culture period, which can clearly distinguish different subclonal cells. The two subclonal cells were analyzed by iCELLigence equipment and to compare with the BHK-21 parental cells.

### Relative cell viability calculation

LS-BHK-21, BHK-21-parental and HS-BHK-21 cells were seeded in 6-well plates 1 day before experiment. At about 60% confluence, the subclones were infected with BEFV (MOI=0.01) and the mock group was incubated with MEM media only. After 48 hours, all experimental groups were trypsinized and counted by hemocytometer under the microscope. Relative cells viability was described by the following equation: relative cells viability = (virus infected group/ mock group) × 100%

### Camptothecin (CPT)-induced cell senescence-like response

Cells (2×10^5^/per well) were seeded in 6 well plates and cultured overnight. CPT were diluted in culture media (250nM final concentration) and added into the cells. After a 24-hour treatment, cells were infected BEFV at an MOI of 0.01. After indicated incubation time, the cell images, virus supernatant, cell lysate or total mRNA were harvested. The virus supernatants were tittered by TCID_50_.

### Western blot analysis for BEFV N protein

The cells were lysed in the RIPA solution (Tools, co. LTD, Taiwan). The protein samples were determined by BCA protein assay (Pierce, Rockford, IL, USA), resolved by SDS-PAGE (20μg/well), and transferred onto a polyvinylidene difluoride membrane (PerkinElmer, MA, USA). After blocking in Hyblock 1min Blocking Buffer (GOAL-BIO, Taiwan) for 1 min at room temperature and washed by PBST (PBS containing 0.05% Tween-20), the membrane was blotted with mouse anti-BEFV N protein antiserum (1:1000) (23). After PBST wash, the membrane was incubated with goat antimouse IgG-HRP (1:5000) antibody. The signal was developed by the ECL plus Western Blot Detection Reagents (GE Healthcare, WI, USA).

### qRT-PCR for senescence related gene expression

Cellular total RNA was harvest and isolated by QIAzol Lysis Reagent (Qiagen, Hilden, Germany). Two micrograms of total RNA was reverse-transcribed with Sensi-FAST™ cDNA Synthesis Kit (Bioline, Memphis, USA). The amounts of p16, p21, IL-6 and β-actin cDNA were determined by QuantiFast SYBR Green PCR Master Mix (Qiagen, Hilden, Germany) at the conditions: 95°C for 3 min and 40 cycles of 95°C for 15 s, 60°C for 30 s and 72°C for 15 s. Primers for each target gene were: p16-F:5’-TCTTGGAAACTCTGGCGATA-3’, p16-R: 5’-GAAGTTACGCCTGCCG-3’, p21-F: 5’-AGTGTGCCGCCGTCTCT-3’, p21-R: 5’-ACACCAGAGT GCAGGACAGC-3’, IL-6-F: 5’-GTCGGAGGTTTGGTTACATA-3’, IL-6-R: 5’-ATCTGGACCCTTTAC-CTCTT-3’ andβ-actin-F: 5’-CCAAGGCCAACCGTGA AAAG-3’,β-actin-R: 5’-TGGCCATCTCTTGCTCGAAGTC-3’. The threshold cycle (CT) value of each target gene was normalized to the β-actin gene, and the mock control group using the 2^-ΔΔCT^ method (24).

### Statistical analysis

Experimental data are expressed as mean ± sd. Statistical analysis of variance between groups was performed by the R Statistics software (4.0.3) with agricolae package. Additional statistical enrichment analysis of GSEA results were performed using the Hypergeometric test to measure the significance. Significant difference of all statistical tests was set at 0.05 (*p* < 0.05).

## Results

### BEFV-infected BHK-21 cells showed morphological changes and upregulated senescence-related gene sets

To study BHK-21 cells’ response to virus infection, experimental infection with BEFV was performed. Infected BHK-21 cells show obvious cytopathic effect (CPE) and cell death. However, some cells survived and became persistently infected, showing enlarged and flattened morphology with apparently more vacuoles inside (**Fig. 1**. in BEFV 48h). These cells can be passaged (**Fig. 1.** in Persistent BEFV-infected BHK-21 (p3)) and variable levels of BEFV can be detected (**Table 1**). To investigate potential gene expression changes within the cells after infection, Gene Set Enrichment Analysis (GSEA) was performed for BHK-21 cells under three conditions: 1) BEFV 24 h.p.i., 2) BEFV 48 h.p.i., and 3) BEFV persistent, when compared with uninfected BHK-21-parental cells. Results showed that senescence-related gene subsets were enriched for all three experimental groups (5, 15 and 9 aging/senescence gene subsets for group 1, 2 and 3, respectively), demonstrating that aging/senescence gene subsets were upregulated in response to virus infection. The regulation of virus infection in cells showed a discontinuous process. Firstly, immune/adhesion-related gene subsets were regulated to respond to virus infection after 24 hours (**Fig. 2**, in BEFV 24 h.p.i.). At 48 h.p.i. and in BEFV-persistent cells, aging/senescence-related gene subsets were upregulated significantly (**Fig. 2**, in BEFV 48 h.p.i. and BEFV persistent). These results suggest that aging/senescence gene subsets upregulation is a potential mechanism for BHK-21 cells to counter BEFV infection.

**Figure 1.**
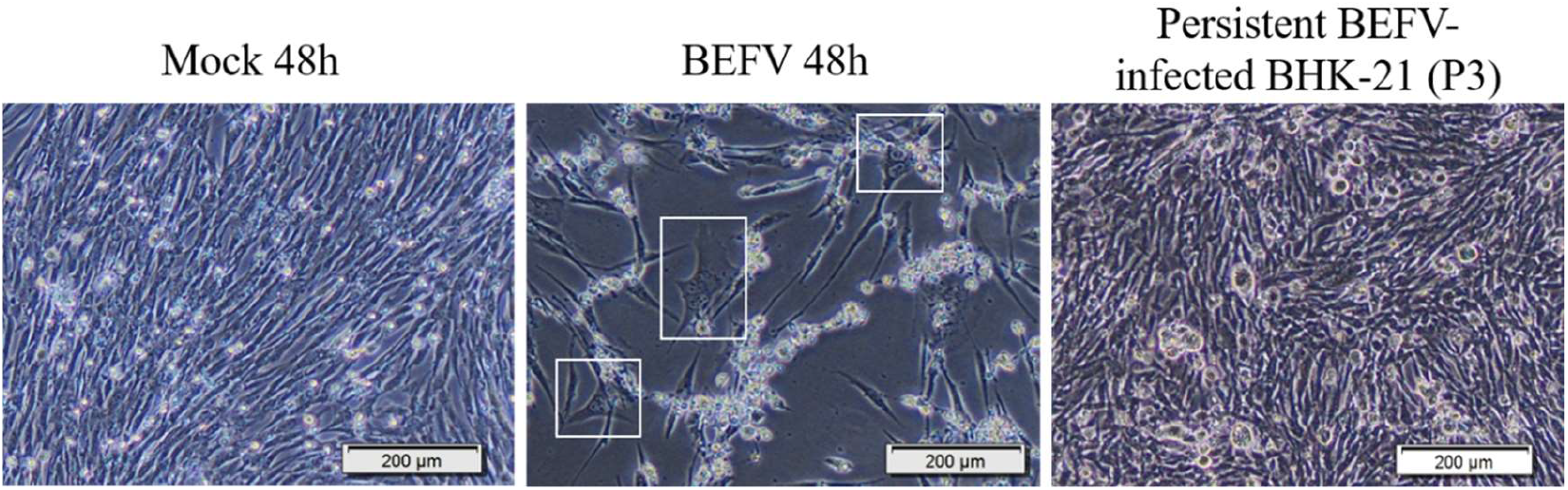
The BHK-21 Cell morphology changes in BEFV infection. Mock 48h: The BHK-21 cells were mock infected for 48h. BEFV 48h: The BHK-21 cells were infected with BEFV for 48h. Persistent BEFV infected BHK-21 cells (p3): The BEFV infected BHK-21 cells were survival and keep culture (passaged for 3 times). The cells were imaged with 400x microscope, and the scale bar showed 200μm.

**Figure 2.**
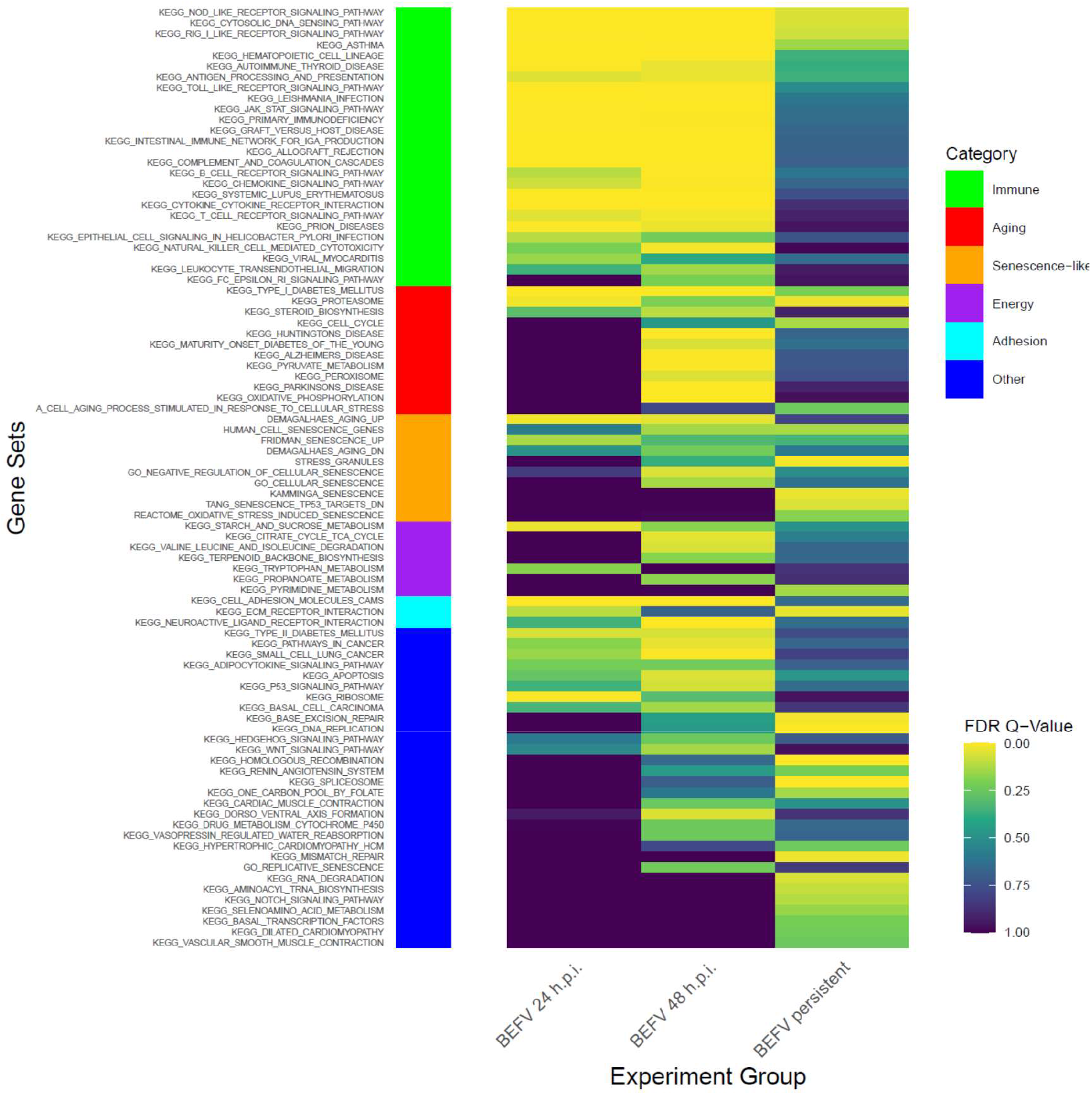
The GSEA results show enriched gene subsets in KEGG pathway with senescence related gene subsets. The results demonstrate FDR value (< 0.25) in KEGG pathway with senescence gene subset at least significance in one experiment group. Side bar in left show categories of gene subsets.

### BHK-21 subclones in senescence-like state are more resistant to BEFV infection

In a slightly different approach, we demonstrated that there are subclones of BHK-21 that show high extent of senescence-like state and are found to be more resistant to BEFV infection. A total of 36 BHK-21 subclones were established from the parental cell line using the limiting dilution method. The subclones were then stained with a senescence-like cellular response marker, SA-β-gal, to determine the extent of senescence-like state (**Supplementary Fig. 1**). Two BHK-21 subclones were picked based on SA-β-gal staining levels and named HS-BHK-21 (for high-staining) and LS-BHK-21 (for low-staining) (**Fig. 3A**). Analysis showed that HS-BHK-21 has higher adhesion ability and larger cell size while LS-BHK-21 resembles the parental BHK-21 (**Fig. 3B**). Upon BEFV infection, HS-BHK-21 showed significantly lower virus titer (85%) when compared to the parental and LS-BHK-21 (**Fig. 3C**), linking higher senescence-like state with resistance to virus infection.

**Figure 3.**
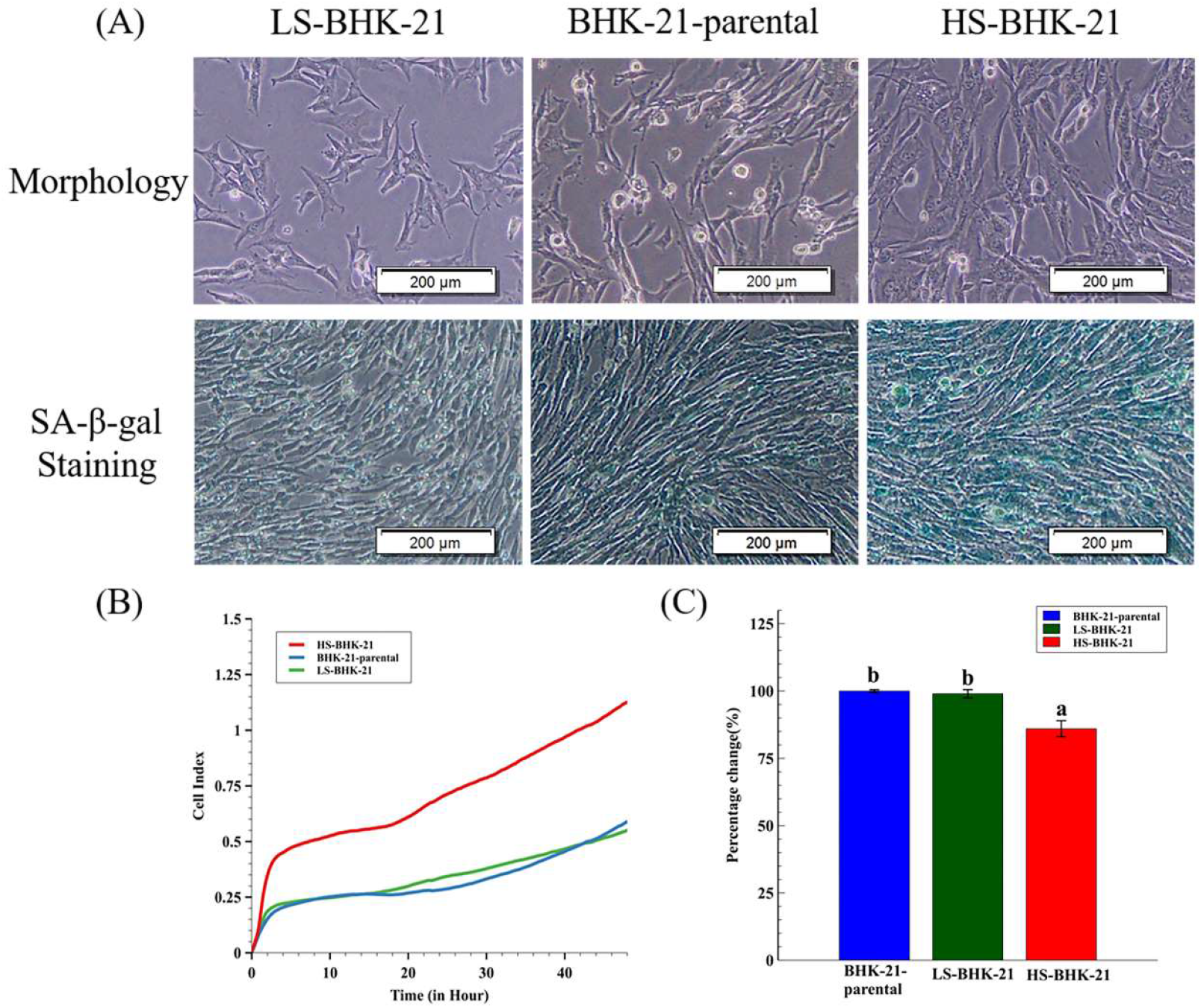
The different SA-β-gal staining property and the virus production of subclonal BHK-21 cells. **(A)** Cell morphology and the SA-β-gal staining results for LS-BHK-21, parental and the HS-BHK-21 cells. The cells were imaged with 400x microscope, and the scale bar shows 200μm. **(B)** The adhesion properties and cell events of the parental and subclonal cells are showed as cell index curve which measured by RTCA iCELLigence™. **(C)** The particular BEFV titer (48 h.p.i.) cultured by the parental and subclonal cells were compared to the BHK-21-parental group and showed as percentage. The means of percentage change with different letters are significantly different (Tukey’s HSD, p < 0.05).

To mimic virus passage during industrial scale virus production, we further investigated how virus titer may change with serial passage of BEFV in the BHK-21 subclones (**Fig. 4A**). Results showed that, for three passages, BEFV titers remained low in HS-BHK-21 when compared to LS-BHK-21 and parental BHK-21 (**Fig. 4B**). This indicates that culturing BEFV in HS-BHK-21 cells may further inhibit virus replication during serial passage. The survival rate of the subclones after each round of virus infection was also tracked (**Fig. 4C**). The three cell populations showed significantly different cell viability after BEFV infection (HS-BHK-21 > BHK-21-parental > LS-BHK-21). HS-BHK-21 showed remarkable resistance to BEFV induced CPE in passage 3. The time-lapse CPE image records in three different senescence states also confirmed the results (https://voutu.be/bVz9VxudRLo). These results indicated that culturing BEFV in HS-BHK-21 cells may further inhibit the virus replication in the following virus passages. A senescence-like cellular function change that may enhance anti-viral response in BHK-21 cells was discovered.

**Figure 4.**
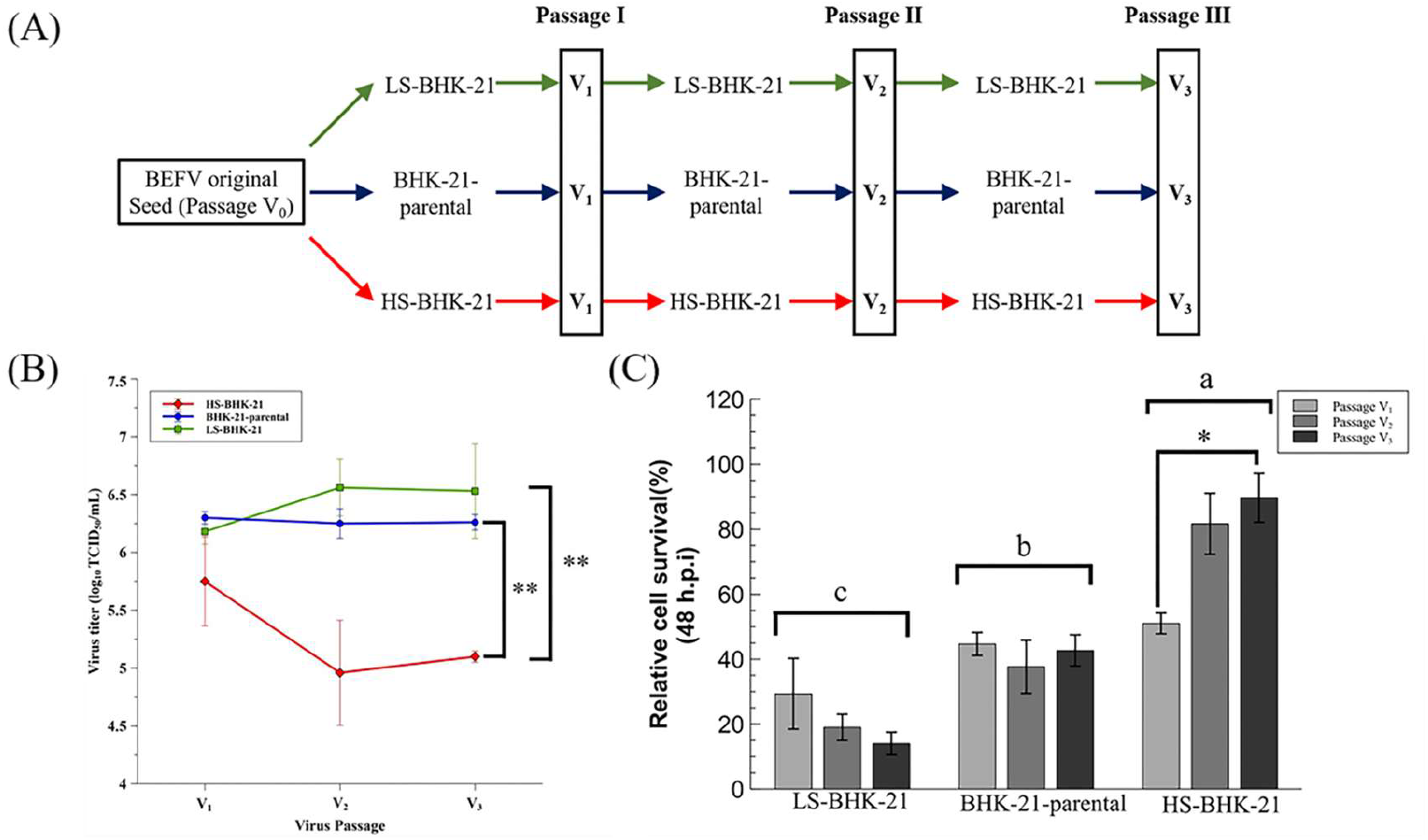
The virus production ability of the parental and subclonal cells in serial virus passages. **(A)** The three serial passages experiment design was showed. **(B)** The virus supernatant of the three continuous serial passages in three different cells population were harvested at 48 h.p.i., and determined by TCID_50_ assay. **(C)** The relative cell survival percentage of the parental and subclonal cells were determined at 48 h.p.i. by directly trypsinized cell counting, and the percentage was compared to the mock infection respectively. Three independent repeats were performed, and were analyzed by one-way ANOVA. Means with different letters are significantly different (Tukey’s HSD, *p* < 0.05). \**p* < 0.05, ** *p* < 0.01 compared to indicated group.

### BHK-21 in drug-induced senescence-like state is more resistant to BEFV infection

Since senescence-like state is drug-inducible, we investigated whether BHK-21 in this drug-induced senescence-like state is more resistant to BEFV infection. The small molecule, camptothecin (CPT), was employed to induce senescence in BHK-21 cells in three senescence state. Upon CPT treatment for 24 h, all cells showed enlarged size and positive SA-β-gal staining (**Fig. 5A upper panel**), giving signs of a senescence-like state. When the treated cells were infected with BEFV, cytopathic effect was significantly inhibited compared to the untreated control (**Fig. 5A lower panel**). A 100-fold decrease in virus titer was observed when compared to the untreated cells (**Fig. 5B**). Also, in the CPT-treated cells, BEFV viral protein N was detected long after infection at 48 h.p.i., verses 24 h.p.i. for the untreated cells (**Fig. 6A**). These results illustrated that BHK-21 in CPT-induced senescence-like state is much less permissible to virus infection. At the gene expression level, further analysis was performed with common senescence-like markers p16, p21, and IL-6 (**Fig. 6B**). Gene expression for p16 was significantly higher in the CPT-pretreated and BEFV 48 h.p.i. groups. For the p12 gene, expression significantly increased in all CPT-treated groups; notably, the expression was even higher for the BEFV 48 h.p.i. group. IL-6 gene expression was significantly higher only after virus infection. Interestingly, p16 and IL-6 gene overexpression was significant in the CPT-pretreated only after BEFV infection for 48 hours. Overall, these results suggest that a senescence-like state was induced by CPT in the BHK-21 cells and upon virus infection, further gene expression change may be observed.

**Figure 5.**
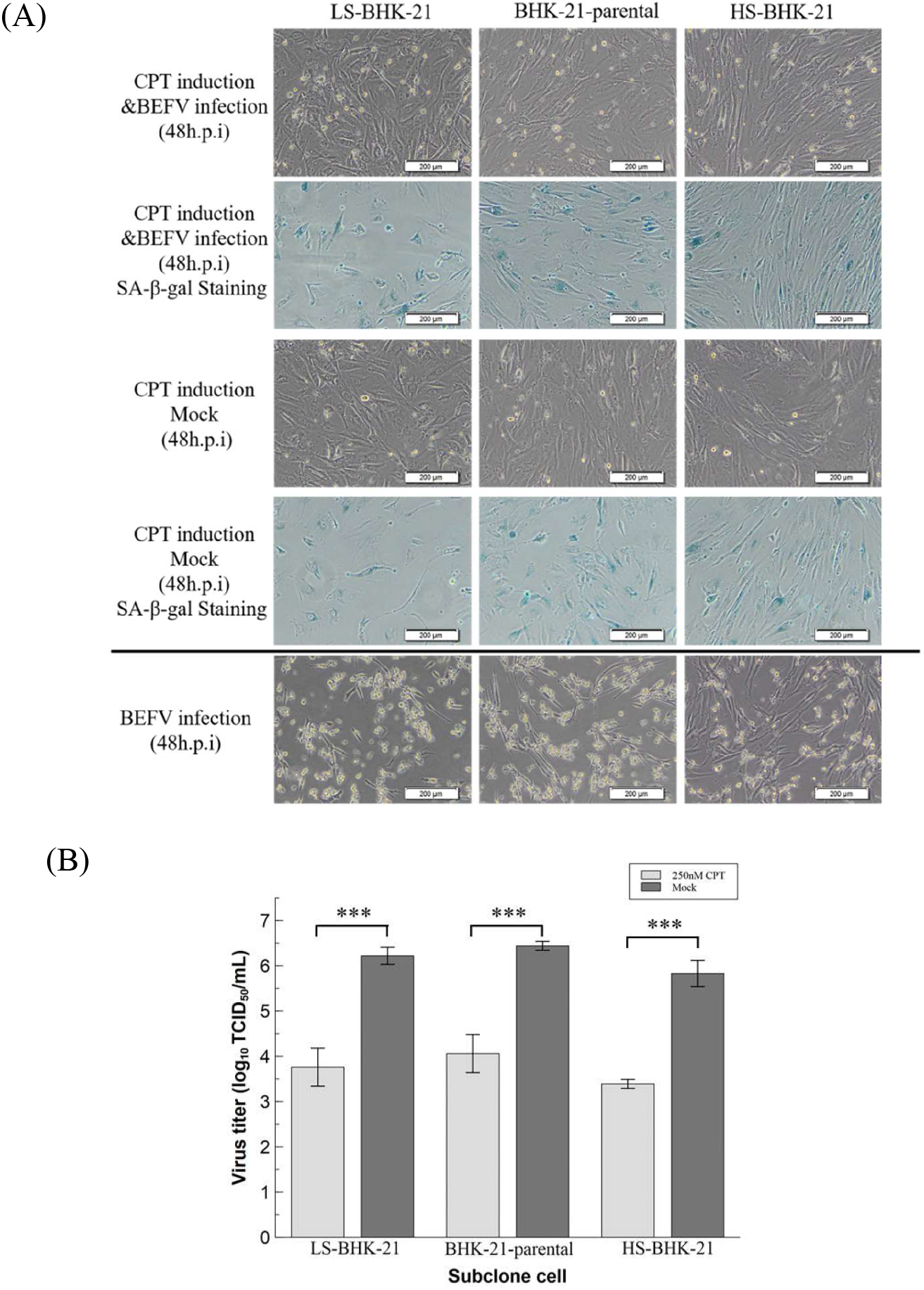
CPT induced senescence-like cell response inhibited virus replication in BHK-21 parental and subclonal cells. **(A)** The parental and subclonal cells were pretreated with CPT (250 nM) for 24hr, and then infected by BEFV or media only as mock control. The senescence-like cell response were determined by SA-β-gal staining and imaged by 400x microscope, and the scale bar shows 200μm. **(B)** The virus titer in the supernatant from CPT pretreatment assay (n=3) were analyzed by TCID_50_ and analyzed by t-test. *** *p* < 0.001

**Figure 6.**
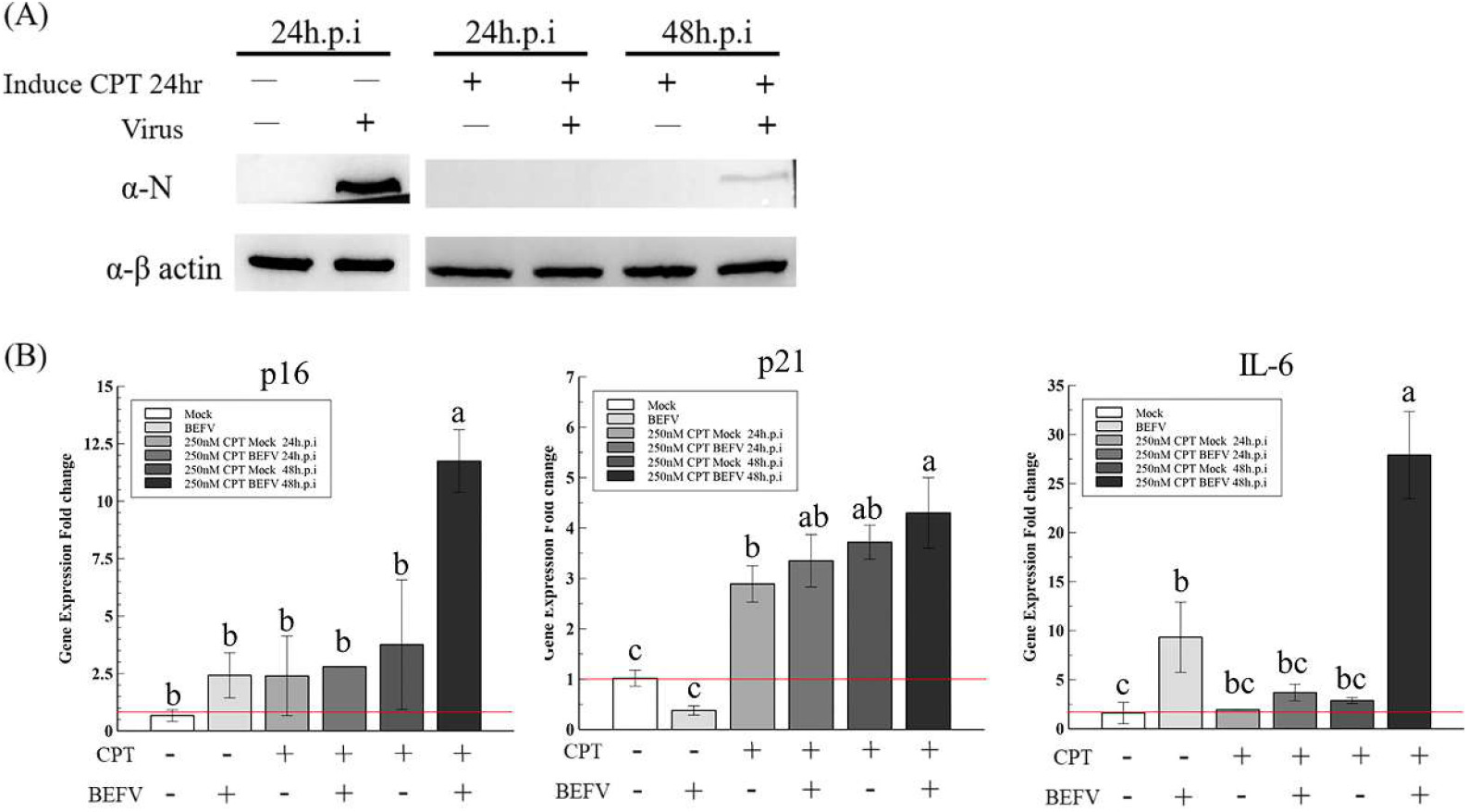
The CPT induced senescence-like response markers, P16, P21 and IL-6, in BHK-21-parental cells with or without BEFV infection. **(A)** The BHK-21-parental cells were pretreated by CPT for 24hr and followed the BEFV infection for 24 and 48hr as indicated. The virus were detected by Western blotting with the mouse anti-N antiserum. **(B)** The cell p16, p21 and IL-6 gene expression in (A) were analyzed by the qRT-PCR respectively and showed as gene expression fold change compare to mock (red line indicate 1 fold). Means with different letters are significantly different (Tukey’s HSD, *p* < 0.05).

## Discussion

In this study, we provided a survey of functional gene sets are affected in BHK-21 upon BEFV infection. Gene set results and cell characteristics together suggested a senescence-like cellular response to viral infection. We further found that subclones of BHK-21 with high senescence marker staining are resistant to BEFV infection. Conversely, BHK-21 subclones with low senescence staining are permissible to virus replication and therefore may be better suited for large-scale bioreactors.

The definition of senescence has been controversial and definitive characterization of the senescent state may be difficult. When first discovered, senescence was associated with DNA damage and telomere shortening. More recent studies found that senescence is a gradually progressing process (25, 26). Senescence-like characteristics may even reverse course before the cell goes into total functional arrest and cell death (27). Furthermore, numerous genes are associated with cell senescence, for example, the p53, p21, p16, and IL-16 (12, 28, 29). In our study, not all markers were examined and induced senescent state may or may not be true senescence, and therefore we used the qualifier ‘senescence-like’ cell response.

During our serial passaging of BEFV in different BHK-21 subclones, we observed a dramatic decrease of virus titer for the HS-BHK-21 subclone, which exhibits high senescence staining. We suspect that the senescence-associated secretory phenotype (SASP) maybe at play. Senescent cells are know to exhibit SASP, which includes the secretion of cytokine, chemokine, extracellular matrix protease, growth factor, and other signaling molecules that can induce senescence in neighboring cells (30). Baz-Martínez M. and colleagues reported that drug-induced senescent A549 cells showed SASP and made daughter cells resistant to vesicular stomatitis virus infection (31). In our study, it is possible that HS-BHK-21 secreted SASP signals into the virus supernatant and resulted in the inhibition of virus replication in the following culture passages. SASP may be the cause of virus titer instability in cell culture systems, especially for high-density bioreactor with a large cell amplifying ratio. There are two ways to solve this problem: the first is the employment of low senescence cells such as the LS-BHK-21 subclone and the second is to adjust the culture media with anti-oxidation or anti-aging components.

## Conclusion

We described a senescence-like cell response in BHK-21 after BEFV infection. BHK-21 subclones showing high senescence staining could inhibit BEFV replication while low senescence-staining subclones are permissive to virus replication. The sub-clones can have utility in virus production for vaccine purposes and can be used for study to develop non-specific anti-viral molecules.

## Declarations

### Consent for publication

Not applicable

### Availability of data and material

The datasets used and analyzed during the current study are available from the corresponding author on reasonable request.

### Authors’ contributions

HYW designed the study. YJZ performed the virus infection assay, protein analysis and gene expression. MKH analyzed the total mRNA expression for the cell functional analysis. CAT isolated the subclonal cells. CYC provided the virus and helped to interpret the data. HCW help to analyze and interpret the data. YJZ, HYW and LTC wrote this paper. All authors have read and approved the manuscript.

## Acknowledgment

We are very thankful to Dr. Li-Ting Cheng in National Pingtung University of Science and Technology for the language editing and the paper reviewing.

